# Spatiotemporal coordination of stem cell behavior following alveolar injury

**DOI:** 10.1101/2022.10.28.514255

**Authors:** Maurizio Chioccioli, Sumner Magruder, John E. McDonough, Jessica Nouws, Tao Yang, David Gonzalez, Lucia Borriello, Brian Traub, Xianjun Ye, Caroline E. Hendry, David Entenberg, Smita Krishnaswamy, Naftali Kaminski, Maor Sauler

## Abstract

Tissue repair requires a highly coordinated cellular response to injury. In the lung, alveolar type 2 (AT2) cells act as stem cells and can replace both themselves and alveolar type 1 cells (AT1); however, the complex orchestration of AT2 stem cell activity following lung injury is poorly understood owing to the inability of tracking individual stem cells and their dynamic behavior over time. Here, we apply live time lapse imaging to ex vivo mouse precision cut lung slice (PCLS) culture and in vivo mouse lung to track individual GFP-labeled AT2 cells following induction of alveolar injury by bleomycin. We observe highly dynamic movement of AT2 cells, including migration within and between alveoli. To map the dynamic evolution of AT2 cell behavior, we introduce Live Cell Encoder (LCE-PHATE), a novel method for converting static snapshots from time lapse imaging into single points representative of entire, dynamic cellular trajectories. Applying LCE-PHATE, we observe the emergence of at least three distinct morphokinetic AT2 cell states associated with AT2 stem cell injury response. Finally, small molecule-based inhibition of Rho-associated protein kinase (ROCK) pathway significantly reduced motility of AT2 stem cells following injury and reduced expression of Krt8, a marker of intermediate progenitor cells. Together, our results uncover motility of alveolar stem cells as a new injury response mechanism in the lung and reveal properties of stem cell motility at high cellular resolution.

## INTRODUCTION

Activation of tissue specific stem cells is a critical response to injury across multiple organs. While many studies have focused on the molecular mechanisms that control stem cell function, relatively little is known about the cellular mechanisms that coordinate their response following injury. This is due to the technical challenges associated with capturing and quantifying individual stem cell behaviors *in situ* in real time, particularly in organs that are difficult to image such as the lung^1–6^.

Some stem/progenitor cells often become motile following injury^7–12^, and this has been associated with their capacity to repair^7–9^. For example, stem cells in hair follicles undergo directed migration to the site of injury in the epidermis, which is critical to their regenerative function^13^. The ability of a stem/progenitor cell to migrate to where it is needed represents an attractive injury response mechanism because it provides a means for localizing repair within a given tissue. However, different tissues have unique structural and microenvironmental properties and tissue-specific response mechanisms associated with the migration of stem cells remain poorly understood.

Using lung alveolar injury as our model, we explore the spatiotemporal coordination of stem cell behavior following administration of bleomycin, a common chemotherapeutic agent known to cause lung injury in mice and humans. Alveoli are located in the distal compartment of the lung and their thin epithelial layer is responsible for gas exchange^14–17^. Upon injury, alveolar type II (AT2) cells function as progenitor cells to support alveolar repair^14,15,18^; however, the spatio-temporal dynamics via which AT2 cells coordinate their response to injury are unknown.

Here, we combine genetic labeling with live time-lapse imaging in vivo and ex vivo to show that AT2 cells become motile following bleomycin-induced lung injury. Tracking hundreds of individual cells over several days, we chart the spatiotemporal evolution of this behavior, demonstrating that motility is an early injury response mechanism adopted by ~30% of all labeled AT2s. We also reveal the unexpected finding that some AT2 cells can exit their native alveolus and navigate through the alveolar pore to adjacent alveoli. Unbiased behavioral mapping reveals the dynamic morphology of motile AT2 cells and identifies diverse behaviors of AT2 cells in response to injury. Mechanistically, inhibition of the Rho kinase pathway significantly reduces the motile phenotype and associated morphological dynamics of AT2 cells, Moreover, inhibition of motility blocks AT2 stem cell fate commitment to a Krt8^+^ phenotype. Together, our findings define motility of AT2 cells as a new injury response mechanism and uncover properties of stem cell motility at high cellular resolution.

## RESULTS

### Time-lapse imaging reveals motility of alveolar stem cells following lung injury

Quantifying the spatiotemporal dynamics of individual alveolar stem cells during injury response requires high-resolution live imaging of large numbers of AT2 cells over an extended period. Previous attempts to image AT2 cells in their native environment have been limited by short durations (<1d) and low throughput^1–6^.To overcome these barriers, we sought to create a platform that would allow for high-resolution imaging of alveolar tissue over a period of days following injury in a controlled and highly reproducible manner. Precision cut lung slices (PCLS) are a well-established ex vivo experimental paradigm in respiratory research^19–21^. Crucially, PCLS retain the cellular complexity and spatial architecture of in vivo alveolar tissue, as well as appropriate cell type ratios, cell-cell and cell-matrix interactions, making them ideal for studying dynamic cell behaviors within the context of their native environment.

To track individual AT2 cells, we crossed Sftpc-CreERT2^22^ with mTmG^23^ mice to generate the SftpcCreER^T2^; mTmG model. In these mice, AT2 lineage cells express GFP while all remaining cells are labeled with tdTomato and appear red. To enable days-long imaging experiments at high resolution, we developed a custom approach based on previously reported PCLS set-ups^24^ with two modifications: i) the addition of a spacer the same thickness as the tissue slice (300 μm) to prevent tissue-distortion and ii) an inlet-opening system to allow media exchange without disturbing the culture (Fig 1a). Our system is compatible with standard confocal microscopy with an incubation chamber to control for temperature, CO_2_ and O_2_.

**Figure 1:**
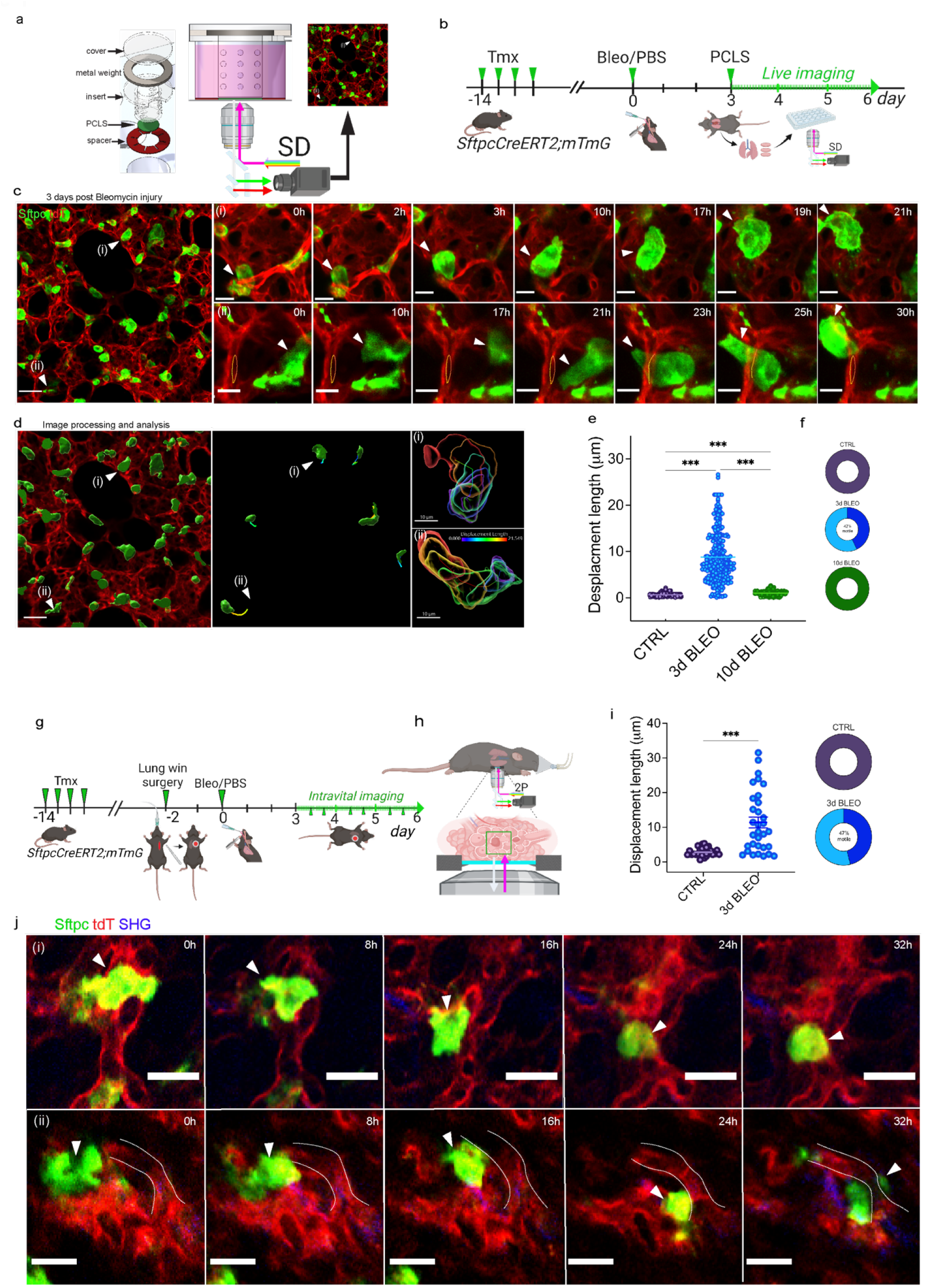
SftpcGFP^+^ AT2 cells become motile in response to alveolar injury. **(a)** Schematic of the ex-vivo imaging system for PCLS **(b)** Schematic experimental workflow for live imaging of *ex vivo* precision cut lung slices (PCLS) generated from injured SftpcCreERT2; mTmG mice. Induction with tamoxifen activates the GFP tag and treatment with bleomycin at day 0 initiates lung injury. Mice are sacrificed on day 3 and lungs harvested to generate 300μm thick PCLS, which are cultured *ex vivo* in a custom-built chamber allowing for continuous imaging for a total of 72 hours. **(c)** Field of view with SftpcGFP-tagged AT2 cells (GFP^+^, green) in situ in mouse alveolar tissue (tdTomato^+^, red) at day 3 post-bleomycin treatment (beginning of imaging). Arrows point to representative cells that exhibit motility over the subsequent imaging period, shown as sequential frames (scale bar 50 μm). (i) AT2 cell (GFP^+^, green) cells moving across an alveolar space (See also Supp Video 1). (ii) AT2 cell (GFP^+^, green) migrating through the alveolar pore (dashed line) into an adjacent alveolus (See also Supp Video 2) (scale bar 10 μm). **(d)** 3D rendering of the same image at the beginning of the imaging experiment shown in **(c)**. Middle panel identifies motile AT2 (GFP^+^) cells at the end of the same imaging experiment and shows cell displacement tracks. Right panel shows separate frames of representative motile cell overlayed and colored as a function of displacement length (i, ii). **(e)** Distribution of displacement length of motile AT2 (GFP^+^, green) cells (violin plots) and total proportion of AT2 (GFP^+^, green) motile cells versus non-motile (donut plots) in control (CTRL dark purple dots) and injured mice at 3 and 10 days post-bleomycin injury (3d BLEO light blue dots). Donut plots were estimated using max displacement length value in control (2.17 μm) as threshold (**e). (g, h)** Experimental approach for intravital imaging of alveolar region in SftpcCreERT2; mTmG. The approach is the same as for the ex-vivo imaging, except that instead of harvesting for PCLS, a lung imaging window is surgically implanted in the mouse for serial imaging rounds across the 72 hour window. **(i)** Displacement length of in-vivo tracked AT2 (GFP^+^) cells in injured mice. **(j)** In vivo imaging through the lung window shows motile AT2 (GFP^+^, green) cells post-bleomycin treated mice (i, ii). Motile AT2 (GFP^+^, green) cells were detected and tracked (See also Supp Videos 4, 5). Statistical significance between conditions was calculated by Kruskal-Wallis Ordinary one-way ANOVA Kruskal-Wallis nonparametric test followed by Dunn’s multiple comparison test; ***p <0.001. Each data point represents a cell. Data are presented as mean ± SEM **(e, i)**.

To capture the dynamic behaviors of AT2 cells post-injury, SftpcCreER^T2^; mTmG mice were treated with Tamoxifen prior to administration of either Bleomycin (treated) or PBS (control) and lungs were harvested 3 days later to obtain PCLS for time-lapse imaging (Fig. 1b). Time-lapse videos revealed a surprising degree of motility of fluorescently-labeled AT2 cells following injury compared to control treated mice. This motility was observed in every video taken in bleomycin treated mice (n = 15 videos; n = 3 mice) (Fig. 1c - f; Suppl. Video 1, 2). To determine the displacement of the motile AT2 cells, we used imaging processing software (IMARIS for Cell Biologist package, Oxford Instruments) to calculate displacement length of individual fluorescent cells in all videos (Fig. 1d). Briefly, individual tracks were generated in three-dimensions for each cell, from which the displacement length was calculated. Overall, we detected movement in approximately 40% of GFP-labeled cells, with notable heterogeneity in the overall displacement of individual cells (Fig. 1e, f). We consider movement as any cell with a displacement value greater than control. On average, cell displacement was 8.5 μm (+/- 0.3277 SEM) over the three days of imaging (days 3 to 6 post-injury), with a range up to ~30 μm (Fig 1e). We repeated our analyses to look at a later time point (days 8 – 10 post-injury) but did not detect any movement (Fig 1e, f, Suppl. Video 3), suggesting that motility of AT2 cells is restricted to the early-phase injury response.

We next sought to validate our findings in vivo. Intravital imaging of mammalian lungs is a formidable challenge in that they reside in a negative pressure chest cavity and undergo constant movement with each breath. Previous attempts to perform intravital imaging of the lung are extremely invasive and allow imaging for only 12 hours or less^1–6^. Recently, a new, minimally invasive window for high-resolution imaging of the lung (WHRIL) was developed that allows repeated intravital imaging at single-cell resolution on exactly the same lung region over a period of days to weeks^25^. We used WHRIL to track individual GFP-labeled AT2 cells during the early injury response in vivo. Briefly, mice underwent surgery to allow implantation of an optically transparent imaging window into the chest wall and allowed to recover for two days prior to treatment with bleomycin to induce lung injury (Fig 1g, h; Supp. Videos 4, 5). Intravital time-lapse imaging began on day 3 post treatment and continued at 2 frames per day imaging window for a total of 3 days. During this time approximately 45% GFP-labeled cells become motile in the bleomycin-treated mice, traversing distances up to ~30 μm, consistent with our ex vivo PCLS imaging system (Fig 1e, i - j). Taken together, our combined *in vivo* and *ex vivo* time-lapse imaging approaches demonstrate that motility is an early-phase response to lung injury of some (but not all) alveolar stem cells.

### Motile alveolar stem cells show a range of dynamic and heterogeneous behaviors

We next sought to explore the dynamic nature of AT2 migration in response to injury. To visualize the relationship between directionality and distance at the single cell level, we plotted cell tracks as spider plots, revealing individual tracks of up to 30μm (Fig 2a, b). These plots also revealed that the individual trajectories of motile cells were non-linear: many tracks appeared to show varying degrees of seemingly exploratory directionality, at times doubling back and/or rapidly changing direction (Fig. 2b; Suppl. Fig 1). Plotting cell acceleration as a function of time revealed a stop-and-start behavior, as opposed to a continuous motion (Fig 2c). We also observed cell division of AT2 cells following injury, both ex vivo and in vivo (Suppl Fig 2; Supp. Videos 6, 7, 8).

**Figure 2.**
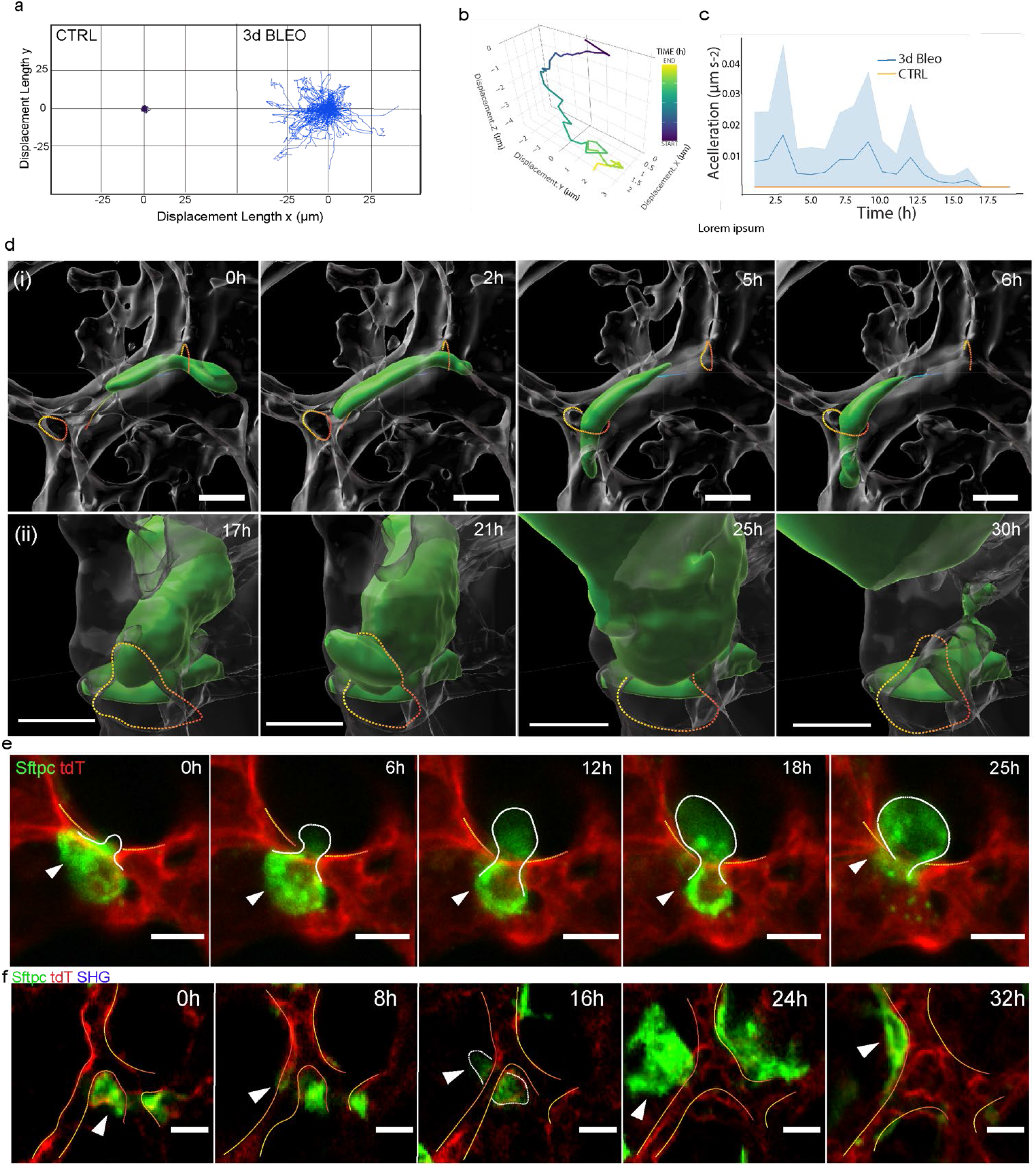
Motile AT2s show a range of dynamic and heterogeneous behaviors. **(a)** Motility tracks of individual cells shown by spider plots of displacement length over time in two dimensions (x, y): saline (CTRL dark purple), 3 days post bleomycin injury (3d BLEO blue tracks); **(b)** example of displacement over time of a single AT2 cell 3 days post-bleo in 3D (x, y, z axis; color map by time). **(c)** average acceleration of AT2s 3 days post-bleomycin (blue) vs AT2s 3 days post saline (orange) over time. (Confidence interval). **(d)** AT2 cell (GFP^+^, green) migrating along the alveolar septum **(i)**. AT2 cell (GFP^+^, green) moving through alveolar pore to the neighbor alveolus (ii) (Supp. Video 2). (Scale bar 10 μm). **(f) (g)** in-vivo AT2 cell (GFP^+^, green) migratory cells move across alveolar boundaries (scale bar 10 μm) (Supp Video 10).

To determine whether AT2 cells can repopulate alveoli distinct from their own, we performed 3D rendering of individual fluorescently-tagged cells and tracked their movement with respect to the surrounding alveolar compartment architecture during the injury response. As expected, we observed dynamic movement of AT2 cells within individual alveoli. We also observed migration of cells between adjacent alveoli with GFP-labeled cells observed moving along the alveolar septum (Fig 2d, i; Supp. Video 9a, 9b) and/or through alveolar pores (Fig 2d, ii; Supp. Video 2, 10, 11). We also observed migration of AT2 cells across alveolar boundaries ex vivo (Fig. 2e), and in vivo (Fig. 2f). Our findings demonstrate that AT2 cells have the capacity to expand their spatial domain during early injury response and can respond to damage either locally (intra-alveolar) as well as between alveoli (inter-alveolar).

### Behavioral mapping reveals morphological parameters associated with motility

So far, our observations have uncovered motility as a dynamic response of AT2 cells to injury and revealed a range of behaviors associated with motile AT2 cells. However, these analyses are qualitative and do not capture nor quantify the full breadth of cell behaviors over the imaging period as well as their relationship to motility. To address this, we developed a novel Live Cell Encoder PHATE (LCE-PHATE) pipeline that converts static snapshots of cellular behavior to low-dimensional embeddings of entire cellular trajectories. Hence, LCE-PHATE is able to quantitatively represent a single cell throughout its entire trajectory as captured by timelapse video.

To accomplish this, LCE-PHATE first encodes high dimensional cellular trajectories containing 68 Imaris features using a sequence-to-sequence (seq2seq) recurrent autoencoder (Supp Figure 3a i, ii). The result is a low dimensional vector embedding representing the trajectory for each cell (Supp Figure 3a iii). We then performed Multiscale PHATE analyses on these vector embeddings (Supp Figure 3a iv). Multiscale PHATE is a method for dimensionality reducing, visualizing and simultaneously clustering data at all levels of granularity^26^. Our Multiscale PHATE groupings organize the heterogeneity of cellular motion over the captured time period. Cells that are close in the embedding undergo similar trajectories in terms of cell type, area, displacement, sphericity, and ellipticity. Altogether, our approach LCE-PHATE pipeline provides two-dimensional visual embeddings that preserve both proximal and distal relationships of the high-dimensional feature space of cellular motility by converting timelapse imaging data of cellular behavior, via a seq2seq autoencoder, to cellular trajectories.

We applied the LCE-PHATE pipeline on 68 individual features exported from videos across 549 GFP-labeled cells from treated (cell n. = 270) versus control (cell n. = 279) mouse lung (Fig 3a). In the resulting visualizations, each point represents a cluster of cells with shared parametric characteristics ascribed by the Multiscale PHATE algorithm (Suppl Fig 3c). Mapping the known control and bleomycin-treated cells onto these data revealed a general separation between the two conditions, as expected. To explore the behaviors that define motility, we examined associations between various features related to cell movement – i.e., displacement, morphology (area, sphericity and ellipticity) – in the AT2 cells (GFP^+^) post-bleomycin injury. We quantitatively mapped each feature onto the ground-truth label and used a color-code to visualize the distribution of values from high to low within individual clusters of cells (Fig 3b - f). Our analyses revealed that cells with a high displacement value (Fig. 3c, top panels, red) – i.e., highly motile cells – are not uniform in their morphology or behavior. For instance, most highly motile cells (Fig. 3c, upper panel, single arrowheads) have correspondingly low surface area values (Fig. 3b lower panel, single arrowheads) except for a small cluster that show both high displacement and high surface area (double arrowheads). With respect to shape, most highly displaced cells tended to be low on the sphericity index and with a higher ellipticity (Fig 3d – f; single arrowheads), though multiple exceptions to this were observed (Fig 3c, double arrowheads). Taken together, our Multiscale PHATE analyses suggest that the motility of AT2 cells following injury proceeds via a range of behaviors, as opposed to a single unified trajectory.

**Figure 3.**
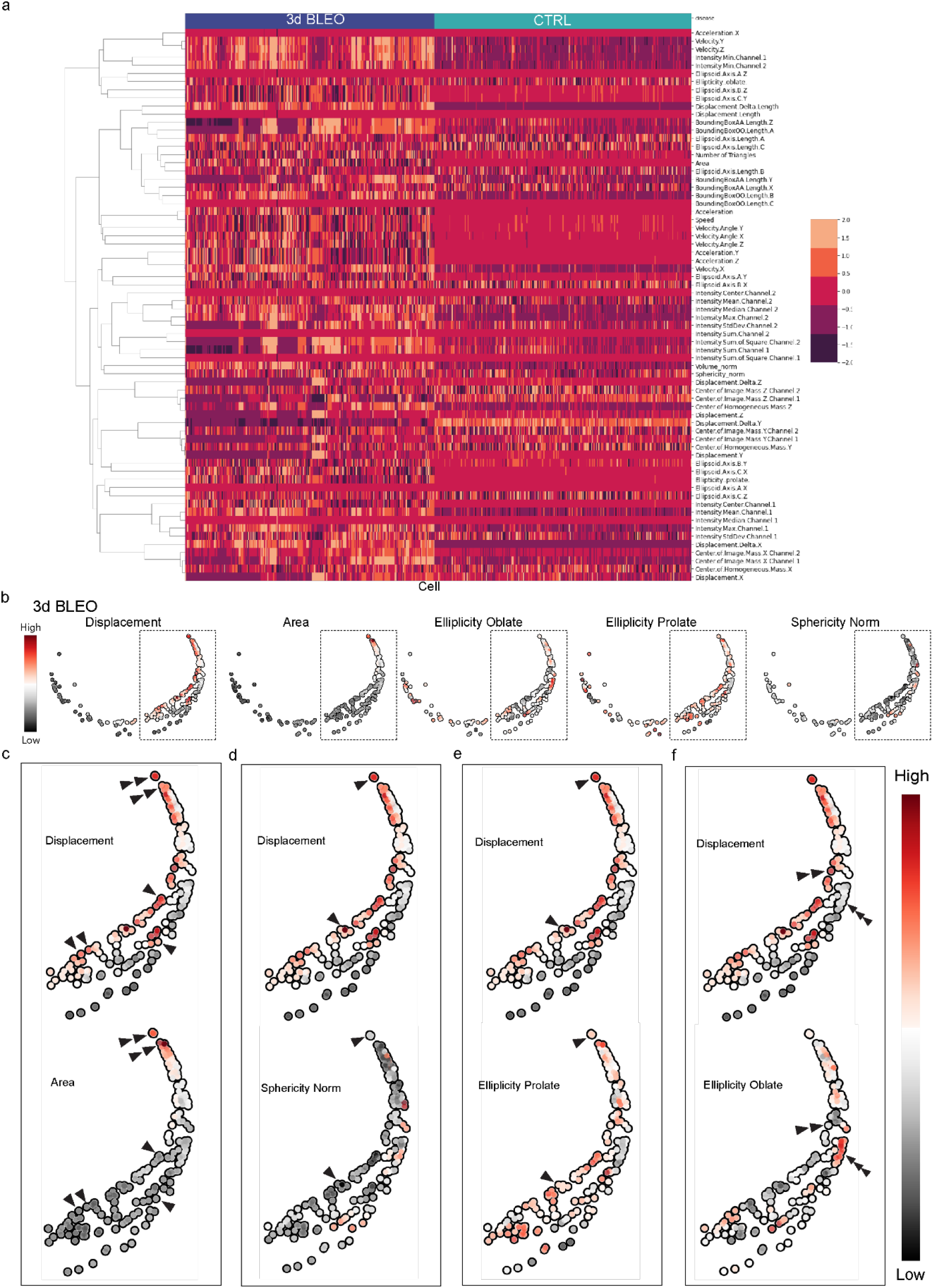
Behavioral mapping reveals morphological parameters associated with motility. **(a)** Heatmap of average Imaris features for bleomycin (purple) and control cells (cyan) using standard scaling. The five features highlighted in multiscale PHATE plots of cellular trajectories **(b)** – Area, Displacement, Ellipticity Oblate, Ellipticity Prolate, and Sphericity – are designated in red font. Recall that each point represents an entire cellular trajectory taken from the latent space of the seq2seq autoencoded. Dendrogram of feature clustering is on the left. **(b)** Multiscale PHATE analyses map of Area, Displacement, Ellipticity Oblate, Ellipticity Prolate, and Sphericity of AT2 cell (GFP^+^, green) 3 days post-bleomycin injury: each feature is mapped onto the ground truth label to visualize the distribution of values from high to low within individual clusters of cells. **(c)** Zoom in of the AT2 cell (GFP^+^, green) showing high displacement and high cell surface area (double arrow heads), and high displacement and low cell surface area (single arrowheads). **(d)** Zoom in of the AT2 cell (GFP^+^, green) showing high displacement but low sphericity (top and middle panels single arrowheads). **(e)** Zoom in of the AT2 cell (GFP^+^, green) showing high displacement and ellipticity prolate (top and middle panels single arrowheads). **(f)** Zoom in of the AT2 cell (GFP^+^) showing high displacement but low ellipticity oblate (double arrowheads), and cells with low displacement length but high ellipticity oblate (triple arrowheads).

### Dynamic behavioral landscape of motile alveolar stem cells

We next sought to validate and further investigate the morphological features identified by LCE-PHATE by returning to our *in vivo* and *ex vivo* datasets (Fig 4). Image analyses revealed that the vast majority of AT2 cells increased in surface area and ellipticity and decreased in sphericity compared to control, ex vivo and in vivo (Fig 4a – d; o, p). However, AT2 surface area and sphericity were poorly correlated with displacement, suggesting no single defining relationship between movement and morphology can explain AT2 stem cell motility, consistent with the conclusion from our LCE-PHATE analyses (Fig 4e, f). On the contrary, while numerous AT2 cells become motile following injury, the behaviors they acquire to do so vary markedly.

**Figure 4:**
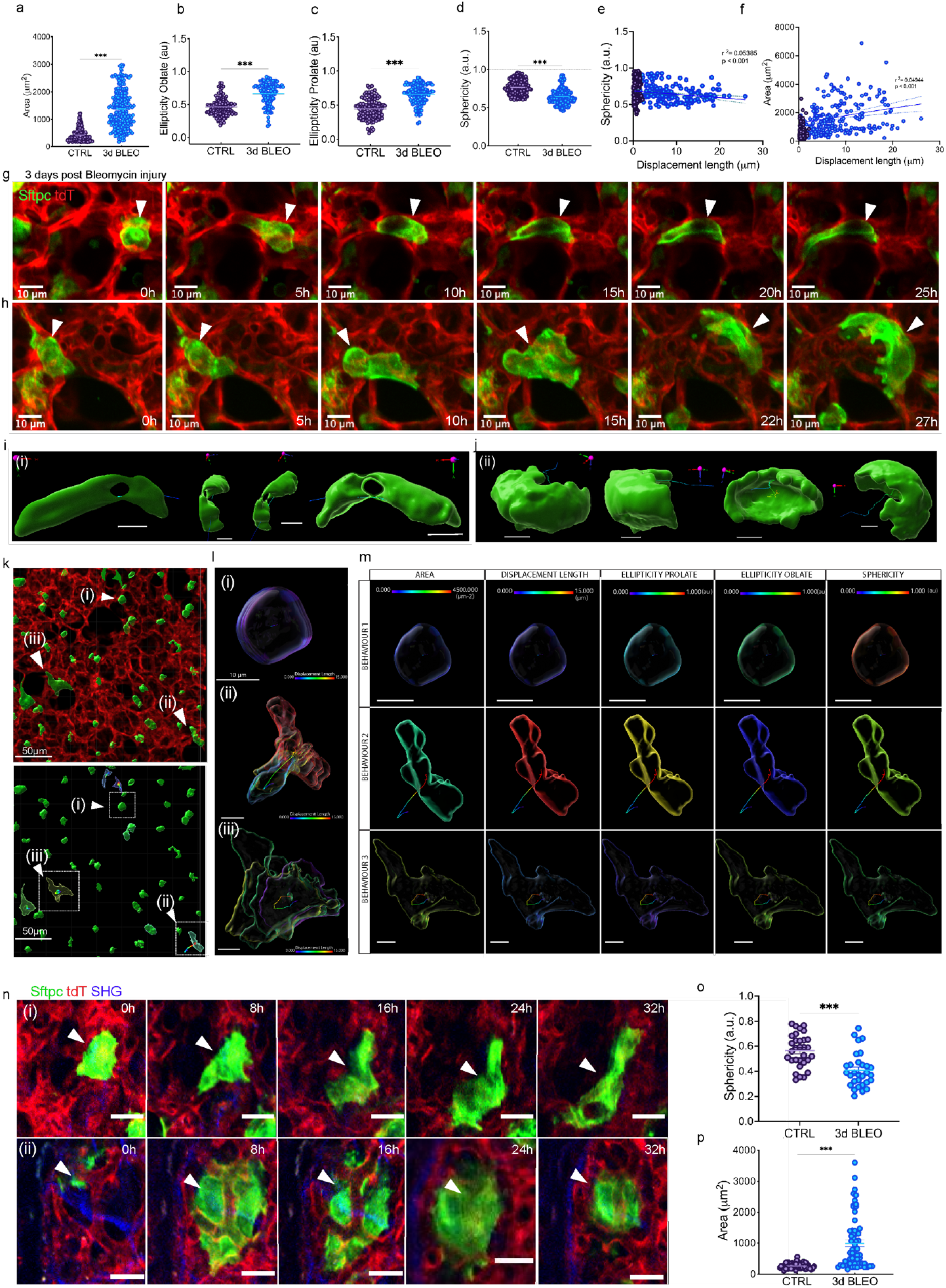
Dynamic behavioral landscape of motile alveolar stem cells. **(a)** Average cell surface area of AT2 cells (GFP^+^) 3 days post-bleomycin injury (3d BLEO light blue dots) and saline treatment (CTRL dark purple dots). **(b)** Average AT2 cells (GFP^+^, green) ellipticity oblate 3 days post-bleomycin injury and saline (CTRL). **(c)** Average AT2 cells (GFP^+^, green) ellipticity prolate 3 days post-bleomycin injury and saline (CTRL). **(d)** Average AT2 cells (GFP^+^, green) sphericity 3 days post-bleomycin injury and saline (CTRL). **(e)** Average AT2 cells (GFP^+^, green) sphericity vs. displacement length (simple linear regression R squared = 0.04944 p<0.001). **(f)** Average AT2 cells (GFP^+^, green) area vs displacement length (simple liner regression R squared = 0.04944 p<0.001). **(g)** AT2 cell (GFP^+^, green) moving and adopting an elongated shape within same alveolus during the movement, **(h)** AT2 cell (GFP^+^, green) adopting a large, flattened, and curved morphology during the movement (ii). 3D renders and displacement tracks of the elongating (i) and enlarging (j) cells from (g) and (h) above. **(k)** Unprocessed and rendered images showing three distinct AT2 cell morphologies. **(l)** overlapping frames of three distinct AT2 cells (GFP^+^, green) phenotypes following injury showing displacement length as a function of color: (i) cuboidal (behavior 1), (ii) elongating (behavior 2), and (iii) enlarging (behavior 3). **(m)** Evolution for each behavioral phenotype spherical (i), elongated (ii), large (iii) according to area, displacement, ellipticity and sphericity as a function of color (max values at the final frame showed). **(n)** in-vivo AT2 cells (GFP^+^, green) showing elongating (i) and enlarging phenotypes (ii). Distribution of sphericity (o) and area (p) in AT2 cells in vivo (Statistical significance between conditions was calculated by Mann-Whitney nonparametric test; ***p <0.001. Each data point represents a cell). Data are presented as mean ± SEM **(a-d)**.

AT2 cells are typically spherical but can become elongated in response to injury^14,15,18,27,28^. We observed this transition in many of our videos of bleomycin treated mice (Fig. 4g, i and Suppl. Vid 12); however, this response was not uniform, as we also observed that AT2 cells adopt a large, flattened, and curved morphology (Fig. 4h, j and Suppl. Videos 13, 14). We further explored parameters associated with these three states: i) spherical ii) elongated and iii) large, flattened (Fig 4j - m). Comparing the parameters of surface area, shape, displacement, and velocity for each behavioral state allowed us to visualize their dynamic behavioral profile and make several observations. First, small, mostly spherical cells are present in resting and injury conditions; these do not typically become motile over time as indicated by very low displacement value (Fig 4m; behavior 1, purple on the color scale). Second, elongated cells – as measured by high ellipticity and low sphericity – undergo much greater displacement over time (Fig 4m; behavior 2). Third, small, mostly spherical cells transform into large, flattened cells, effectively increase in surface area and size in situ over time without displacement (Fig 4m; behavior 3). We also observed these motile behaviors in vivo (Fig. 4n; Suppl. Video 15 (enlarging), 16 (elongating)). Together, these results suggest that AT2 cells respond to injury via at least three behaviorally distinct paths.

### Inhibition of motility prevents alveolar stem cell fate commitment

We next sought to investigate the potential physiological relevance of AT2 stem cell motility injury responses using small molecule inhibitors of cellular migration. PCLS from bleomycin-treated mice were treated with either Y-27623 dihydrochloride, a small molecule inhibitor of the Rho-associated protein kinase pathway that regulates cell adhesion and migration, or saline solution. We first performed continuous live imaging to track changes in cell behavior and morphological parameters in response to Y-27623 (Fig. 5a). Upon treatment, we observed a dramatic reduction in displacement of GFP cells, indicating a near-total abrogation of cell movement (Fig. 5b). We also observed an absence of all morphological parameters previously identified and associated with the motile AT2 cells (Fig 5c – g). Together, these results demonstrate that Y-27623 treatment inhibits AT2 cell motility and abrogates the behavioral phenotypes associated with motile AT2 cells.

**Figure 5:**
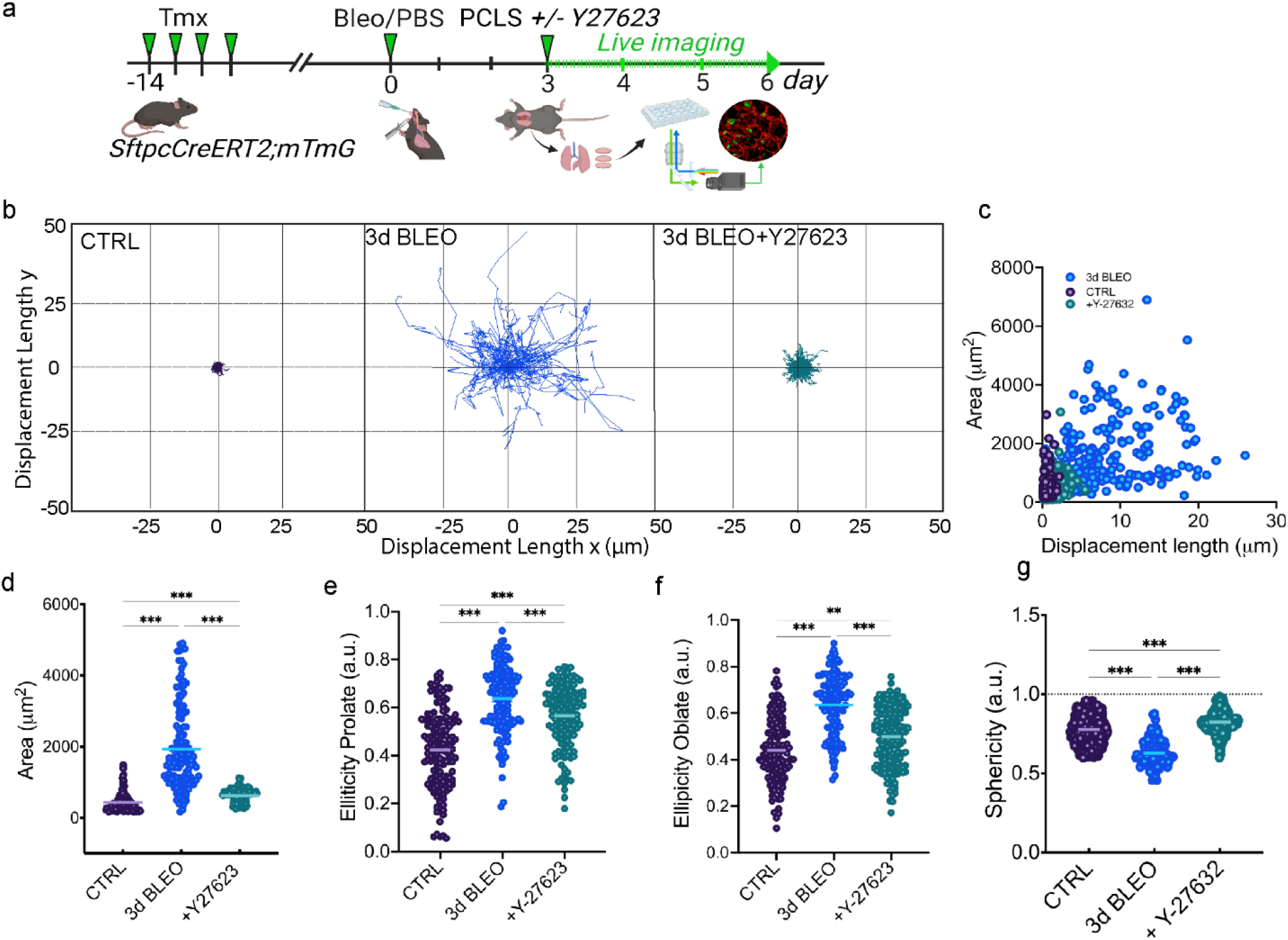
Y-27632 reduce AT2 motility post injury. **(a)** Schematic experimental workflow for live imaging of ex vivo precision cut lung slices (PCLS) generated from injured SftpcCreERT2; mTmG mice (1.5 U kg^−1^ bleomycin), and control treated mice (vehicle). Mice are sacrificed on day 3 post injury and harvested to generate 300 μm thick PCLS. PCLS are cultured with/without Y-27632 ROCK inhibitor (20 μM) containing media (3d BLEO, 3d BLEO + Y-27632) or vehicle (CTRL). **(b)** Displacement length over time as spider plots of individual cell tracks 3 days post-saline (CTRL dark purple tracks), 3 days post-bleomycin injury (3d BLEO blue tracks) and 3 days post-bleomycin injury + Y-27632 ROCK inhibitor (20 μM) (3d BLEO + Y-27632 green tracks). **(c)** AT2 cells (GFP^+^, green) area vs. displacement length; **(d)** AT2 cells (GFP^+^, green) area. **(e)** Ellipticity prolate, oblate (**f**) and sphericity **(g)**. Statistical significance between conditions was calculated by Kruskal-Wallis Ordinary one-way ANOVA Kruskal-Wallis nonparametric test followed by Dunn’s multiple comparison test; ***p <0.001. Each data point represents a cell. Data are presented as mean ± SEM **(d-g)**.

Following injury, AT2 cells are responsible for generating new cells in order to repopulate damaged alveoli^14,27,29,30^. Recent studies have revealed that one of the earliest steps of AT2 cell differentiation is their commitment to a Krt8^+^ transitional cell state^28,31–33^ and that this transitional cell state is enriched for the expression of motility genes^31,34^. To better understand the relationship between the onset of stem cell motility and cell fate commitment we analyzed publicly available single-cell sequencing data of AT2 cells after injury (GSE141259)^31^ using Manifold Interpolating Optimal-Transport Flows for Trajectory Inference (MIOFLOW). Using MIOFLOW, we were able to interpolate continuous gene expression dynamics from time course single-cell sequencing data. We found a continuous increase in the expression of multiple motility related genes across the first 7 days as AT2 cells transition to Krt8^+^ intermediate progenitor cells (Fig. 6a).

**Figure 6:**
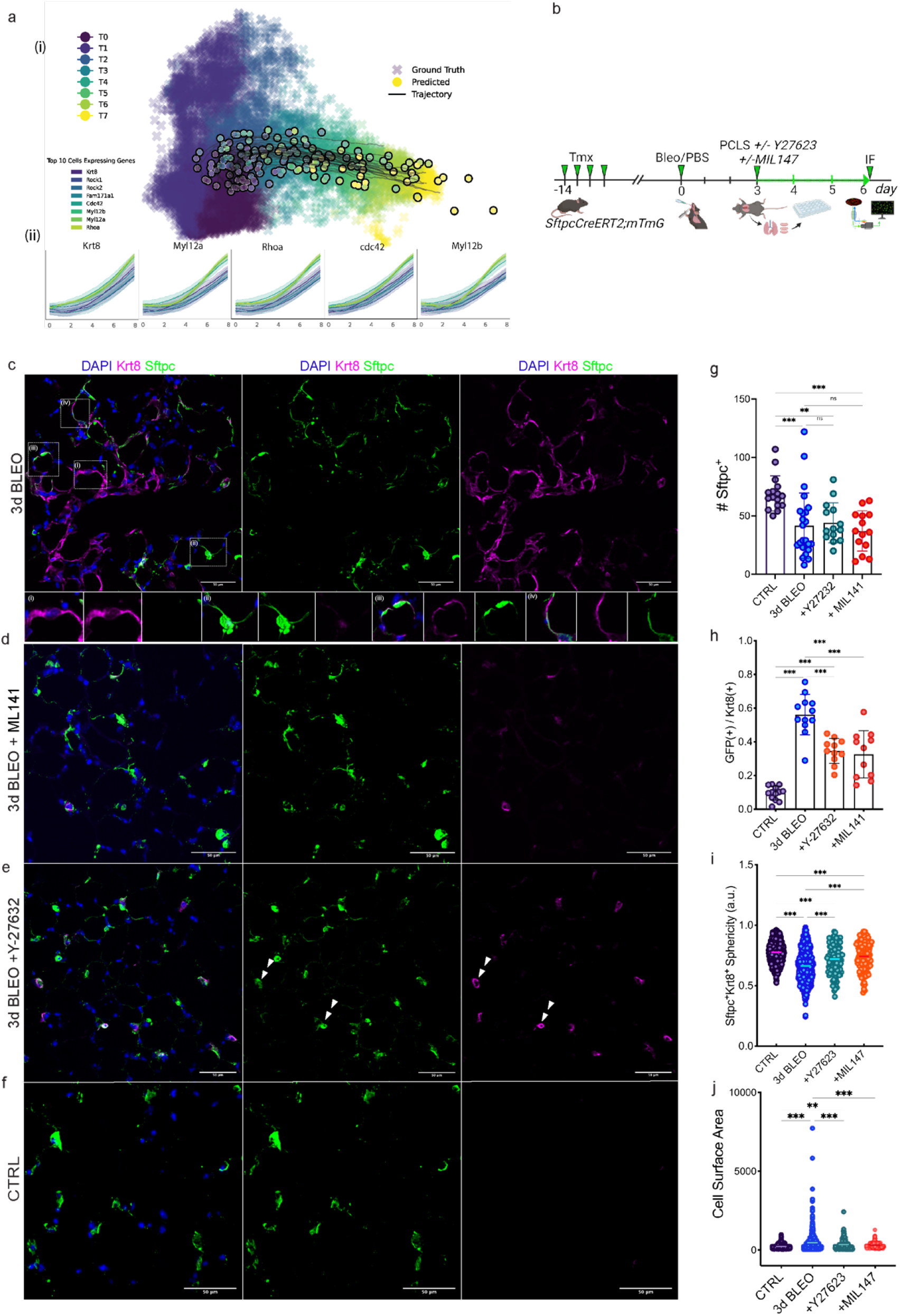
inhibition of motility reduces the number of Krt8^+^ cells in alveolar regions following injury. **(a)** MIOFlow trajectory analysis of AT2 single-cell RNA-seq results over first 7 days following treatment with bleomycin: MIOFlow trajectory analysis of AT2 single-cell RNA-seq results over first 7 days following treatment with bleomycin. (i) MIOFLow trajectories in the 100 dimensional PCA embedded space (first two dimensions shown). Points are marked by solid circles. Cell trajectories are marked by black lines. (ii) Gene trends of cell trajectories over pseudotime. For the designated genes the top 10 cells expressing these cells (indicated by color) are shown for other genes of interest (the band represents confidence interval). **(b)** Schematic experimental workflow: ex vivo precision cut lung slices (PCLS) generated from injured SftpcCreERT2; mTmG mice (1.5 U kg^−1^ bleomycin), and control treated mice (vehicle). Mice are sacrificed on day 3 post injury and lungs harvested to generate 300 μm thick PCLS. PCLS are cultured with/without Y-27632 (20 μM) containing media, and ML141 (3d BLEO, 3d BLEO + Y-27632, 3d BLEO + ML141) or vehicle (CTRL). At day 6 post injury PCLS are fixed, stained, and imaged. **(c, d, e, f)** Immunofluorescence staining of PCLS shows nuclei DAPI (blue), Krt8 (magenta), GFP (green) (Scale bar: 100 μm) for control (bottom panel) and injury (top panel) and injury treated ex vivo with either Y-27632 or ML141 (middle panels). **(g)** GFP^+^ Krt8^+^ double positive cells plotted as a fraction of all GFP^+^ cells for each condition. **(h)** total number of GFP^+^ cells for conditions. **(i)** GFP^+^Krt8^+^ cells sphericity, and area **(j)**. Number and shape of positive cells were quantified by cell surface software analysis (IMARIS, Oxford Instruments) on 15 randomly chosen FOVs (FOV= 620 × 620 μm; n =3 mice/conditions). Statistical significance between conditions was calculated by Kruskal-Wallis Ordinary one-way ANOVA Kruskal-Wallis nonparametric test followed by Dunn’s multiple comparison test; ***p <0.001. Each data point represents a cell. Data are presented as mean ± SEM **(f-i)**.

To understand the functional relationship between AT2 cell motility and Krt8^+^ fate commitment, we sought to determine the effect of inhibition on the generation Krt8^+^ cells. We repeated the above experiment using Y-27623 to inhibit AT2 cell movement as well as ML141 to selectively inhibit Ccd42/Rac1, another Rho-family GTPase^35–38,40^ (Fig 6b). Following three days of treatment with either the inhibitors or control (saline), we stained for Krt8 expression and quantified the number of GFP^+^ cells as well as the number of Krt8^+^ cells per field of view (n = 15 fovs) across all conditions (Fig 6c-h). We observed no significant difference in the number of GFP^+^ AT2 cells between injury and injury plus treatment with either Y-27623 or ML141 inhibitors on the (Fig 6g). In contrast, the number of Krt8^+^ cells per fov dramatically decreased following blockage of motility in the injury condition with either inhibitor (Fig 6c – e, right column).

These results show that inhibition of motility reduces the number of Krt8^+^ cells in alveolar regions following injury, however, they cannot determine whether those Krt8^+^ cells arose from AT2 cells or from elsewhere. To address this, we used imaging analyses to calculate the number of Krt8/GFP double positive cells per fov and plotted these as a fraction of all GFP^+^ cells (Fig 5), allowing us to consider Krt8 expression only in *Sftpc*:GFP^+^ AT2 cells. We observed a significant reduction in the fraction of Krt8^+^/GFP^+^ cells per fov following inhibition of motility after injury. This was associated with a phenotypic shift towards the profile described as behavior 1 above (non-motile cells), with increased sphericity and decreased surface area (Fig. 6h, i). For the fraction of Krt8^+^/GFP^+^ cells still present following inhibition (Fig 6e, double arrowheads), we speculate that these were committed to the Krt8^+^ fate prior to inhibition, and so escaped the blockade. Taken together, our results reveal that small molecule inhibition of the Rho kinase pathway with either Y-27623 or ML141 rapidly reduces AT2 motility and associated behavioral profiles, resulting in fewer Krt8^+^ expressing AT2 cells. These results suggest an important link between AT2 cell motility and cell fate commitment.

## DISCUSSION

Our study defines the spatiotemporal coordination of AT2 cells following alveolar injury and identifies a link between motility as a cellular mechanism and stem cell fate commitment. We show that motility of AT2 cells is restricted to the early post-injury phase and demonstrate the emergence of three morphokinetically distinct states of AT2 cells in response to injury. Motile AT2 cells can respond to damage both within their own alveolus (intra-alveolar) as well as between alveoli (inter-alveolar) via intra-alveolar pores, revealing an under-appreciated role for alveolar pores in the communication between alveoli in tissue homeostasis. Blockage of motility leads to a marked reduction of Krt8 expression in AT2 cells, suggesting their motility may be linked to cell fate commitment to an intermediate state. Together, these results open up a new avenue for investigating the cellular mechanisms that regulate stem cell motility and how these, in turn, are linked to tissue repair.

One intriguing observation is the range of behaviors adopted by motile AT2 cells following injury. We did not detect a single, stereotypical morphological trajectory as cells become motile. Rather, we found that motility proceeds via multiple distinct and dynamic behaviors. We hypothesize that this could be due to heterogeneity in the AT2 microenvironment as the injury response proceeds. Differences in tissue architecture pose unique and fundamental challenges for proper stem cell function; hence, it could be that AT2 cells are sensing changes in their microenvironment and adapting their behaviors accordingly. Whether this sensing proceeds via mechanosensing, growth-factor based signaling or a combination of both remains to be explored.

In this study we observed the movement of AT2 cells through alveolar pores and between adjacent alveoli. Such pores have previously been reported as pores of Kohn and hypothesized to be integral for maintaining pressure equilibrium among alveoli ^14,39,41–43^. Our observation that AT2 cells can be triggered to exit their native alveolus and transit to a new alveolus raises the intriguing question of how this process is regulated. Given the close association of motile AT2 cells with the alveolar septum and their exploratory behavior, mechanosensing may be involved, particularly given the changes in mechanical pressure that cells experience in damaged alveoli^44^.

One of the advantages of continuous imaging is that it provides the opportunity to directly observe the dynamic evolution of complex processes and the cellular mechanisms that govern them. In the future, intersecting live imaging approaches described here with genetic perturbations could help to dissect not only the molecular pathways that regulate AT2 stem cell motility, but also the consequences when they are perturbed. This approach has been used successfully in other injury/repair contexts where live imaging is well-established ^45–48^. We expect that a similar approach could help to uncover key regulators of AT2 motility following injury and dissect the molecular pathways that drive them toward alternate behaviors.

### Limitations of the study

The limitations of this study revolve around imaging: primarily the limited time-span (days) as well as the trade-off between the high-resolution, large dataset generated by our ex vivo PCLS platform and the lower resolution data generated by in vivo imaging. The difference in resolution was due to the light scattering due to the heterogeneous nature of the lung tissue. Nonetheless, using a combined approach, we were able to provide strong evidence of AT2 cell motility following injury ex vivo and in vivo and then turn to PCLS for detailed morphokinetic analyses at high resolution across hundreds of cells. Such a large dataset is required for multiscale analyses to identify the behavioral “rules” of AT2 stem cell motility in an unbiased and quantitative manner.

Although PCLS culture is viable for over to a week, our imaging system did not allow for continuous live imaging for more than 3 days^21^. This is limiting, since alveolar repair proceeds over a much longer period^14,15,27^ and hence we are unable to investigate either the dynamic behavior of AT2 cells over this timespan, or the ultimate effect on repair of blocking their motility. Further improvements in the PCLS chamber design may help to overcome this limitation.

## ACKNOWLEDGMENTS

We thank Dr Valentina Greco (Yale Genetics) for thoughtful discussion and feedback. We thank Dr Derek Toomre and Felix Rivera Molina (Yale Cell Biology Cellular Imaging using New Microscopy Approaches, CINEMA LAB), for imaging equipment, technical support, and fruitful discussions.

## AUTHOR CONTRIBUTIONS

Conceptualization, M.C., C.H., S.M, S.K., N.K., M.S.; Methodology, M.C., S.M., S.K., D.E.; Software, S.M., S.K., D.E. X.Y.; Formal Analysis, S.M., S.K., J.M., X.Y., D.E., D.G.; Investigation, M.C., L.B., B.T., T.Y., J.N.; Resources, N.K., M.S., S.K., D.E.; Data Curation, M.C., J.M., S.K., X.Y., D.E.; Writing – Original Draft, M.C., C.H.; Writing – Review & Editing: M.C., C.H., S.M., M.S., S.K., D.E., N.K.; Supervision, M.C., S.K., D.E., M.S., N.K.; Project Administration, M.C., S.K., D.E.; Funding Acquisition, S.K., M.S., N.K., D.E.

## FIGURES TITLES AND LEGENDS

## STAR METHODS

### Lead contact

Requests for further information or reagents may be addressed to the lead contact, Maurizio Chioccioli

### Materials availability

This study did not generate new unique reagents.

### Data and code availability

Any additional information required to reanalyze the data reported in this paper is available from the lead contact upon request.

## EXPERIMENTAL MODEL AND SUBJECT DETAILS

### Mice

SftpcCreERT2; mTmG was generated by crossing *Sftpc*^*tm1(cre/ERT2)Blh*^*(Sftpc-CreER*^*T2*^) (stock number 028054, Jackson Laboratory) with *B6*.*129(Cg)-Gt(ROSA)26Sor*^*tm4(ACTB-tdTomato,-EGFP)Luo*^*/J (mTmG)* (stock number 007576, Jackson Laboratory). All animal husbandry and experiments were approved by the Institutional Animal Care and Use Committees (IACUC # 07867) at Yale University and Einstein College of Medicine (Protocol# 00001079) and in accordance with U.S. Government Principles for the Utilization and Care of Animals Used in Research, Teaching and Testing. Mice were provided clean bedding, *ad libitum* water and food, and nestlets. Mice were maintained on a C57BL/6 background and genotyped to determine their control versus appropriate lineage tracing genotypes and allocated accordingly at weaning. Both sexes were used for this study.

## METHOD DETAILS

### Lineage tracing

For all lineage tracing experiments, 4 doses of tamoxifen (0.1mg/g body weight in corn oil) (Sigma Aldrich) were administered intraperitoneally 14 days prior to bleomycin (McKesson 1129996) treatment. For bleomycin injury, mice were administered (1.5U/kg dissolved in 50 μL PBS or 0 U/kg in 50 μL PBS for control) of bleomycin intratracheally under isoflurane anesthesia and then monitored daily.

### Precision cut lung slices (PCLS) preparation

Precision cut lung slices were generated from mouse lung as previously described^49^. The procedure was performed under sterile conditions. A shielded I. V. catheter (BD Insyte Autoguard 381423) was carefully inserted through the trachea up to a millimeter above the bifurcation of the principal bronchi and fixed in place by suture. After cannulation and dissection of the diaphragm, lungs were flushed via the heart with sterile phosphate-buffered sodium chloride solution. Using a syringe pump, approximately 1mlL of 37°C 1.5% by weight low-melting-point agarose (Sigma Aldrich, A9414) in sterile DMEM/Ham’s F12 cultivation medium (Gibco, 12634010), supplemented with 100 U/ml penicillin, 100 μg/ml streptomycin, and 2.5 μg/ml amphotericin B (Sigma Aldrich, A2942), was slowly injected to fully inflate both lungs. The trachea was ligated with thread to retain the agarose inside the lungs. Afterwards, the lungs were excised and transferred to a tube with cultivation medium and cooled on ice for 10 min to allow the agarose to gel. Each lung lobe was separated and cut with a vibratome (Compresstome VF-300-0Z; Precisionary Instruments) to a thickness of 300 μm using a speed of 10–12 μm·s^−1^, a frequency of 80 Hz, and an amplitude of 1 mm. Agarose was largely washed out upon slicing. PCLS were obtained from the middle 2/3rds of the lobe (to ensure similar-sized slices) and placed in a 24-well plate (1 PCLS/well) filled with 600 μL cultivation medium / well. The slices were allowed to float in the media throughout the culture period. PCLSs were cultivated at 37°C in humidified conditions containing 5% (volume/volume) CO_2_ in 24-well plates under submerged conditions with changes of medium every 2h for the first 12h to remove excess agarose from the tissue, and every other day.

### Live imaging of PCLS

Time-lapse imaging on PCLS was conducted as previously described^24^ with some modifications. After 24h in culture conditions, PCLS were transferred at the center of the well of an uncoated 24 well glass bottom plate with #1.5 cover glass (Cellvis P241.5HN) (1 PCLS/well). Each well was provisioned with a metal spacer of the same thickness as the tissue slice (300 μm) to keep samples in place and prevent tissue-distortion. PCLSs were kept immobilized and flat on the glass bottom by placing a 12 mm diameter porous membrane (0.4 μm) inlet-opening system (BrandTech 782821) on top of the PCLS and weighing it down with 1.66 g flat metal washer (Fig 1b). To ensure and maintain optimal culture conditions, each PCLS was submerged in 800 μL of phenol red-free DMEM F-12 (supplemented with 100 U/mL penicillin, 100 μg/mL streptomycin, and 2.5 μg/mL amphotericin B). PCLS were then left in the incubator for 2 h to allow the samples to acclimate and settle prior to image acquisition. The specific design of the cylinder insert membrane enabled a proper nutrient exchange to the PCLS. Wells were covered with removable plastic lids to minimize evaporation. The 24-well plate was then transferred to a pre-equilibrated and humidified stage-top environmental control chamber (OKO LAB H301-NIKON-TI-S-ER), where temperature (37ºC), humidity (95-100%), and gas concentration (5% CO_2_) were monitored and maintained through. Time-lapsed fluorescence imaging (1 frame/h for 72h, 5 × 1020 × 1020 μm randomly selected field of view / PCLS), were performed on PCLSs (n = 6 / conditions), generated from SftpcCreERT2; mTmG mice (n = 5 each condition) 3 days post bleomycin injury or saline as control, using a micro-lens enhanced dual spinning disk confocal microscope equipped with an sCMOS Camera (ANDOR Zyla 5.5) and using a 20X PlanApo 0.75NA objective (Nikon).

### Surgical protocol for implantation of the WHRIL

Briefly, 4 doses of tamoxifen (0.1mg/g body weight in corn oil) (Sigma Aldrich) were administered intraperitoneally 14 days prior to the experiment. Two days prior experiments, mice (n = 2 for each condition), underwent surgery to allow implantation of an optically transparent imaging window into the chest wall and allowed to recover for two days prior to treatments. The surgery for permanent implantation of a thoracic optical window designed to enable serial imaging of the murine lung via high-resolution multiphoton microscopy was conducted as previously described^25^. For bleomycin injury, mice were administered (1.5U/kg dissolved in 50 μL PBS; 0 U/Kg in 50 μL PBS for control) of bleomycin (McKesson 1129996) intratracheally under isoflurane anesthesia and monitored daily. All procedures have been performed in accordance with guidelines and regulations for the use of vertebrate animals, including prior approval by the Albert Einstein College of Medicine Institutional Animal Care and Use Committee.

### Intravital imaging

Intravital imaging was performed as previously described^28^. Briefly, on day 3 post treatments mice were anesthetized using 5% isoflurane and placed on the microscope stage. An environmental enclosure maintained the mouse at physiological temperatures, and vitals were monitored using pulse oximeter (MouseStat for PhysioSuite, Kent Scientific). Anesthesia was maintained at 0.75-1.5% isoflurane.

Imaging was performed on a custom-built, two-laser multiphoton^50^ microscope. Imaging began on day 3 post treatment. Settings: About 10 z-stacks of FOVs / mice (340×340 μm, 11 z-slices spanning 30 μm from the lung surface) were chosen and recorded twice a day for a total of 3 days. In vivo microcartography was used to re-localize a region of interest between each session over the 3 days as previously described^28^. All images were captured in 16 bit using a 25x 1.05 NA objective lens and acquired with two frame averages.

### Intravital imaging processing

For each FOV, images at each time point were coarsely registered to each other, manually. This was accomplished by identifying the same vascular features at each timepoint and translating (in x, y, and z) and rotating (in x & y) each z-stack to the position of the features in the first time point. Further fine registration was accomplished by application of the HyperStackReg plugin^51^, a registration algorithm for 5D data sets based upon the StackReg plugin^52^. For each FOV, two cells were selected for area, circularity, and motility analyses. A threshold value for quantifying the cell area was first chosen manually in Fiji (ImageJ 1.53b). Then, a custom written ImageJ script was used to automatically extract the cell area and circularity data based on the threshold value. It also saved the cell ROIs for later motility analysis. Cell motility was quantified using the ROI Tracker plugin in ImageJ (ImageJ 1.52b). The final cell displacement data was extracted from the ImageJ plugin generated tracks in MATLAB (MathWorks, Natick, MA).

### PCLS Immunofluorescence staining

Paraffin embedded blocks of PCLS were deparaffinized in xylene and decreasing concentrations of ethanol in distilled water. They were then placed in citrate pH-6 epitope retrieval buffer at 95°C for 20 min, then cooled and permeabilized in PBS with 0.1% tween and 0.2% triton for 5 minutes, and subsequently rinsed and washed in PBS with 0.1% tween. Slides were incubated in CAS-Block (Life Technologies, #008120) for 10 minutes. The primary antibodies (applied overnight at 4ºC) were rabbit anti-SPC (1:200; Millipore AB3786), rat anti-KRT-8 (1:100, DSHB TROMA-1), and Goat anti-GFP (1:200; Rockland 620010115M). Slides were washed with TBS with 0.1% tween and then incubated for 1 hour at room temperature with the secondary antibodies donkey anti-rabbit AF-647 (Invitrogen A32795), donkey anti-rat AF-488 (Invitrogen A21208), and donkey anti-goat AF-555 (Invitrogen A21432). Slides were washed with PBS with 0.1% tween and coverslips were mounted using anti-fade mounting media with DAPI (Vectashield H-1200-10). Images were captured using a micro-lens enhanced dual spinning disk confocal microscope equipped with a sCMOS Camera (ANDOR Zyla 5.5) using a 20x PlanApo 0.75NA objective (Nikon).

### Inhibition of motility

Ex vivo precision cut lung slices (PCLS) generated from injured SftpcCreERT2; mTmG mice (1.5 U kg^−1^ bleomycin), and control treated mice (vehicle) were sacrificed on day 3 post injury and lungs harvested to generate 300 μm thick PCLS. PCLS were cultured with/without Y-27632 dihydrochloride (20 μM) (TOCRIS 1254) containing media, and ML141 (Calbiochem 217708) (3d BLEO, 3d BLEO + Y-27632, 3d BLEO + ML141) or vehicle (CTRL). At day 6 post injury PCLS are fixed, stained, and imaged.

### Live Cell Encoder PHATE (LCE-PHATE) Pipeline

The cells we chose to track with the IMARIS imaging software were exported. Notably, the exported data is not normalized with respect to the viewing window. Therefore, we first normalized the IMARIS features with respect to each tracked cell of the experimental triplicate groups and aggregated all of the cells into a cohesive dataset of 549 cells (270 bleomycin treated cells and 279 control cells). We then employ recurrent neural networks (RNN) to create a sequence-to sequence (seq2seq)^53^ autoencoder using gated-recurrent units (GRU)^54^ for both the encoder and decoder, as GRUs empirically demonstrate higher specificity^55^ than long short-term memory (LSTM) units. In essence, the GRU is given the capacity to forget via the reset gate. Generally, a RNN takes a given input sequence (*x*_1_, …, *x*_*T*_) and produces the corresponding output sequence (*y*_1_, …, *y*_*T*_) via the repeated application of the equation:

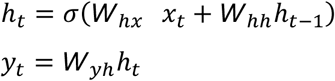

The GRU tweaks this slightly with the aforementioned reset gate 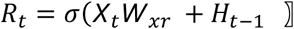 and update gate *Z* _*t*_ = σ(*X*_*t*_ *W*_*xz*_ + *H*_*t* −1_ *W*_*hz*_ + *b*_*z*_)as well as update the hidden state 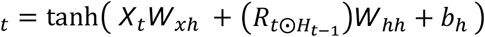.

The input sequence (*x*_1_, …, *x*_*T*_) corresponds to the IMARIS features of a given cell at time points *t* = 1, …, *t* = *T* and the output sequence 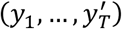 is identical to the input. Further we leverage the GRU hidden state to reduce the dimension of the feature vector *x*_*i*_. We refer to this latent space as the trajectory embedding for a given cell *c*_*i*_. Our GRU seq2seq autoencoder was trained using the Adam optimizer and an L1 loss. Use of the autoencoded *trajectory embedding* is key as it takes the feature matrix of static time-lapse imaging of cells and produces - via the hidden state of the GRU from the encoder - a single feature vector representing the entire cellular activity.

Following the conversion from a static series of images to a singular representation of the dynamic process, we further analyze cellular behavior by employing the Potential of Heat-diffusion for Affinity-based Trajectory Embedding (PHATE) algorithm. PHATE constructs a diffusion geometry of the trajectory embedding via the local similarity between cells through the following process: compute Euclidean distances on the trajectory embedding of cells, applying the alpha-decaying kernel with adaptive bandwidth to produce affinities, instantiating the diffusion process via a Markovian random-walk, converting these to potential distances by taking the negative log, and then reducing the dimensionality via non-metric multidimensional scaling (MDS). Following the processing on the trajectory embedding from the seq2seq autoencoder, yields a low dimensional embedding with one point per cell that respects local and global distances of the cellular behavior from the original IMARIS imaging software export. While this itself is novel and highly informative, we further innovate by applying multiscale PHATE (MSPHATE) which leverages diffusion condensation to visualize and cluster cellular behaviors of the *entire* time-lapse imaging process at fine and coarse grain resolutions (see Supp figure 4a).

From the seq2seq autoencoder to convert varying duration snapshots sequences of cellular behavior into single representative trajectory embeddings per cell to MSPHATE’s coarse and fine grain visualization and clustering of the cellular trajectories, the LCE-PHATE pipeline enables differentiation between behaviors rather than sole datapoints.

### MSPHATE analysis

We performed Multiscale PHATE26 analyzes on the trajectory latent space produced by a seq2seq autoencoder with python 3.9 and the Pytorch package (1.11.0), on the data generated from the time-apse videos by IMARIS Cell Biology Software (Oxford Instruments) as part of the LCE-PHATE pipeline. MSPHATE produces coarse to fine grain resolutions of cellular behaviors by applying diffusion condensation^56^. If matrix *P* are the transition probabilities on our dataset, then powering*P* i.e. *P*^*t*^ produces the transition probabilities after taking *t*-steps. Further note that the diffusion operator *P* applied to a vector averages the values of the vector over a small neighborhood. Thus the iterative diffusion condensation process is represented by *f*_*k*_^(*t* +1)^ = *P*_*t*_*f*_*k*_^(*t*)^ = *P*_*t*_ *P*_*t* −1_ …*P*_1_*P*_0_*f*_*k*_, *t* > = 0, where *f =* (*f*_1_, …, *f*_*n*_) are coordinate functions and *f*_*k*_ ∈ ℝ^*n*^ is the vector of a given coordinate function applied on the data and the *P* ^*t*^ uses diffusion smoothing to contract the data. Then the diffusion operator is recomputed on the contracted (or condensed data) and reapplied. Therefore, the dataset *X* = (*x*_1_, …, *x*_*n*_) iteratively becomes coarse grained as the coordinate functions *f*_*k*_ of the condensation process at time *t* + 1 are derived through the repeated application of the diffusion process *P*^(*t*)^ = (*P*_*t*_, …, *P*_*0*_). Where application of diffusion once results in the following description of the data, *<u>X</u>* = {*<u>x</u>*_1_,…, *<u>x</u>*_*N*_} = {(*Pf*1(*z*1),…, *Pfn*(*z*1)), …, (*Pf*1(*zN*),…, *Pfn*(*zN*))}, where the data is modeled by *xi* = *f*(*zi*) for some *zi* ∈ *M*^*d*^ and *M*^*d*^ is the manifold. This process of applying the diffusion operator onto the original coordinate functions *fk* smoothes them adaptively removing high frequency noise pulling points towards the local barycenters, where *X*(*t* + 1) is derived through the previously described process, i.e. 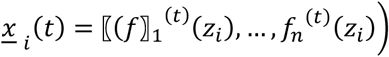, to produce *f*_*k*_ ^(*t+1*)^. MSPHATE applies this process by building the MSPHATE operator and then fitting the provided data with said operator to construct a diffusion condensation tree. The coarse grained coordinates stem from merging clusters when at a given time step their pairwise distances are less than some threshold.

### MIOFLOW scRNAseq data analysis

The integrated dataset after the preprocessing was subset for time less than or equal to 9 days and for cells labeled as either AT1, AT2, or Repair cells. The PHATE algorithm was employed to produce a low dimensional embedding which respects local and global high dimensional distances. We then applied Slingshot^61^ algorithm to generate pseudotime labels in the low dimensional space along the principal curve. The first 100 principal components (PCs) of the Strunz et al., 2020 scRNA-seq dataset (GSE141259)^31^ generated as part of the PHATE algorithm, were then labeled with pseudotime and used as the input for Manifold Interpolating Optimal-Transport Flows for Trajectory Inference (MIOFlow) model. Analysis was limited specifically to alveolar epithelial cells. As a neural ordinary differential equation (Neural ODE), MIOFlow learns continuous population dynamics from the static time samples of the Strunz et al., 2020 data. MIOFlow’s geodesic autoencoder enabled us to invert from the embedded space back to gene space producing the gene trends seen in figure 7. Gene trends for motility genes as indicated by Gene Ontology are selected for analysis, across which we see general increasing expression over pseudotime.

### QUANTIFICATION AND STATISTICAL ANALYSIS

Statistical methods relevant to each figure are outlined in the figure legend. Statistical analysis was performed using GraphPrism 9 (GraphPrism Software) and R (v4.2.1). Statistical methods relevant to each figure are outlined in the figure legend. Data were considered significant if p < 0.05. Mice were randomly assigned to treatment or control groups, while ensuring inclusion criteria based on gender and age. The number of animals shown in each figure is indicated in the legends as n = x mice per group. Data shown are presented as mean ± SEM.

## SUPPLEMENTAL INFORMATION TITLES AND LEGENDS

**Supp Figure 1**. Ex-vivo 3D tracks of AT2 cells (GFP^+^) displacement length in x, y and z axes (μm) over time of AT2 GFP^+^ green 3 days post-bleomycin injury (color map by time).

**Supp Figure 2. Cell proliferation and migration** of “a” (AT2 GFP^+^ green) into “b” ex-vivo **(a)** and in-vivo **(b)** 3 days post-bleomycin injury (scale bar 10 m).

**Supp Figure 3. Live Cell Encoder PHATE Pipeline for generating multiscale PHATE plots. (a)** Cell feature matrix: Individual features exported from Imaris imaging software are aggregated into a cohesive matrix with each cell, at a given point in time, represented by a row and each feature (e.g., area, displacement, etc.) as a column. There are a total of 139 Bleo cells, 279 Ctrl cells, and 188 Rock cells. **(ii)** Cell2Cell (or Seq2Seq) autoencoder: a latent trajectory space embedding is learnt for each cell by using a traditional Seq2Seq autoencoder model e.g., the encoder takes the sequence of features for a *single* cell over time and attempts to reconstruct this cellular behavior (as indicated by features) with the decoder. iii.) The cells trajectory feature matrix - the latent space encoding of each cell - is extracted from the Cell2Cell AutoEncoder. Whereas in (i) each time point for each cell is designated by a given row, now (iii) one row represents a cell over all observed time. iv.) The trajectory feature matrix is fed into multiscale PHATE for unsupervised analyses. **(b)** Multiscale PHATE resolutions. Here the learnt PHATE embedding of Bleo cells can be viewed at different visual resolutions. On the left each cell’s trajectory embedding is represented by a single point. Decreasing resolution towards the right, cells are aggregated where; larger dots indicate more cells have been merged. While other embedding methods are popular (e.g. UMAP, tSNE), PHATE was designed specifically to preserve both local and global distances. Further the multiscale facet of PHATE assists with visual reasoning tasks. **(c)** Multiscale PHATE feature projection. In this panel different cellular conditions are indicated by the y position in the panel (3d Bleo, 3d Bleo vs. Ctrl, and 3d Bleo v 3d BLEO+Y27623, (see Figure 3). The mean feature value a given a cell over time is projected into the embedding space as indicated by the colorbar. Comparative visual analysis shows cell specific features. For example, the second row (3d Bleo vs. Ctrl) first (ground truth labels) and third (displacement) columns show that Bleo cells have higher displacement on average.

