## Supplementary Figures and Videos for "Spatiotemporal coordination of stem cell behavior following alveolar injury"

### SUPPLEMENTAL INFORMATION

### Supp Figure 1

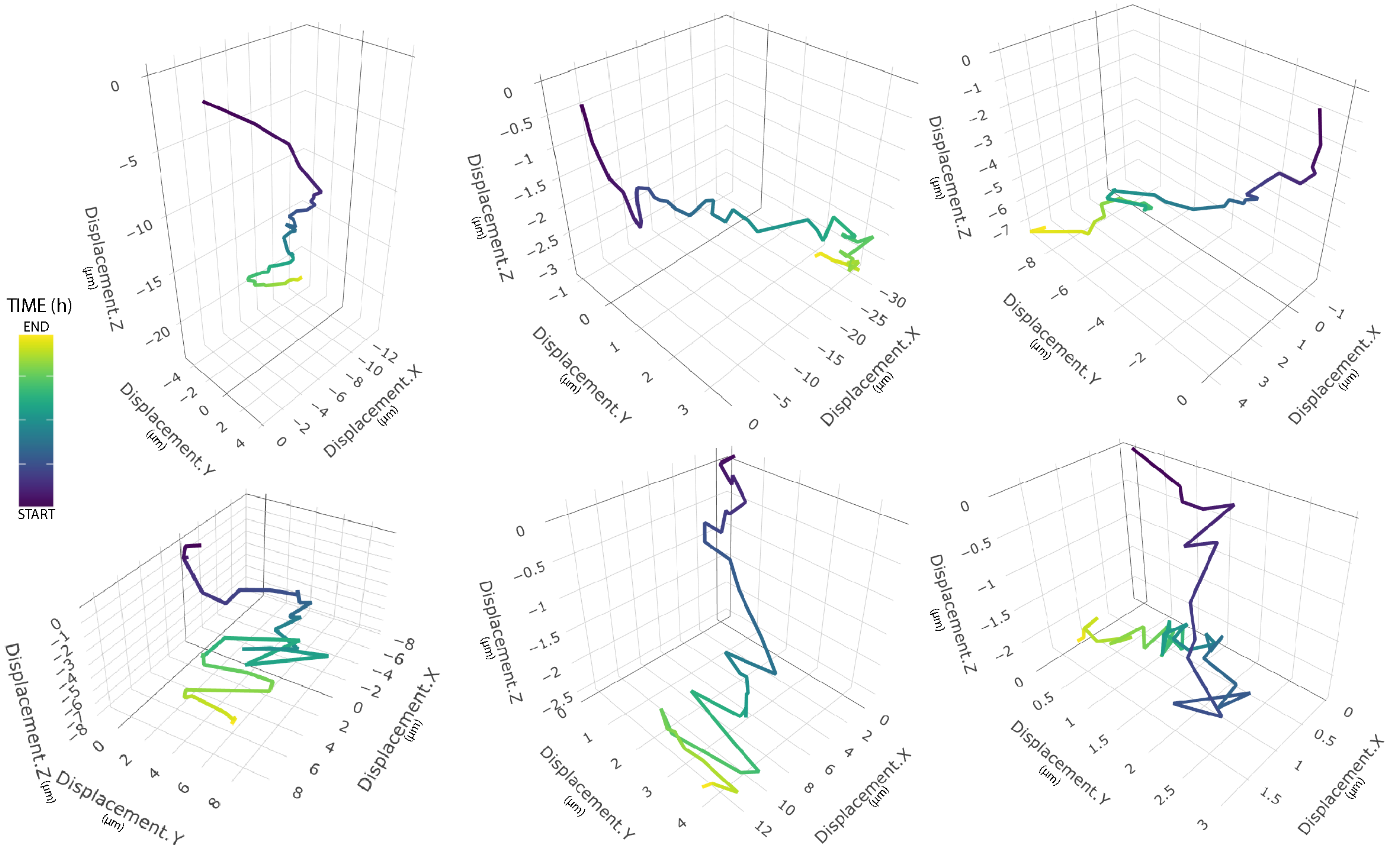

### Supp Figure 2

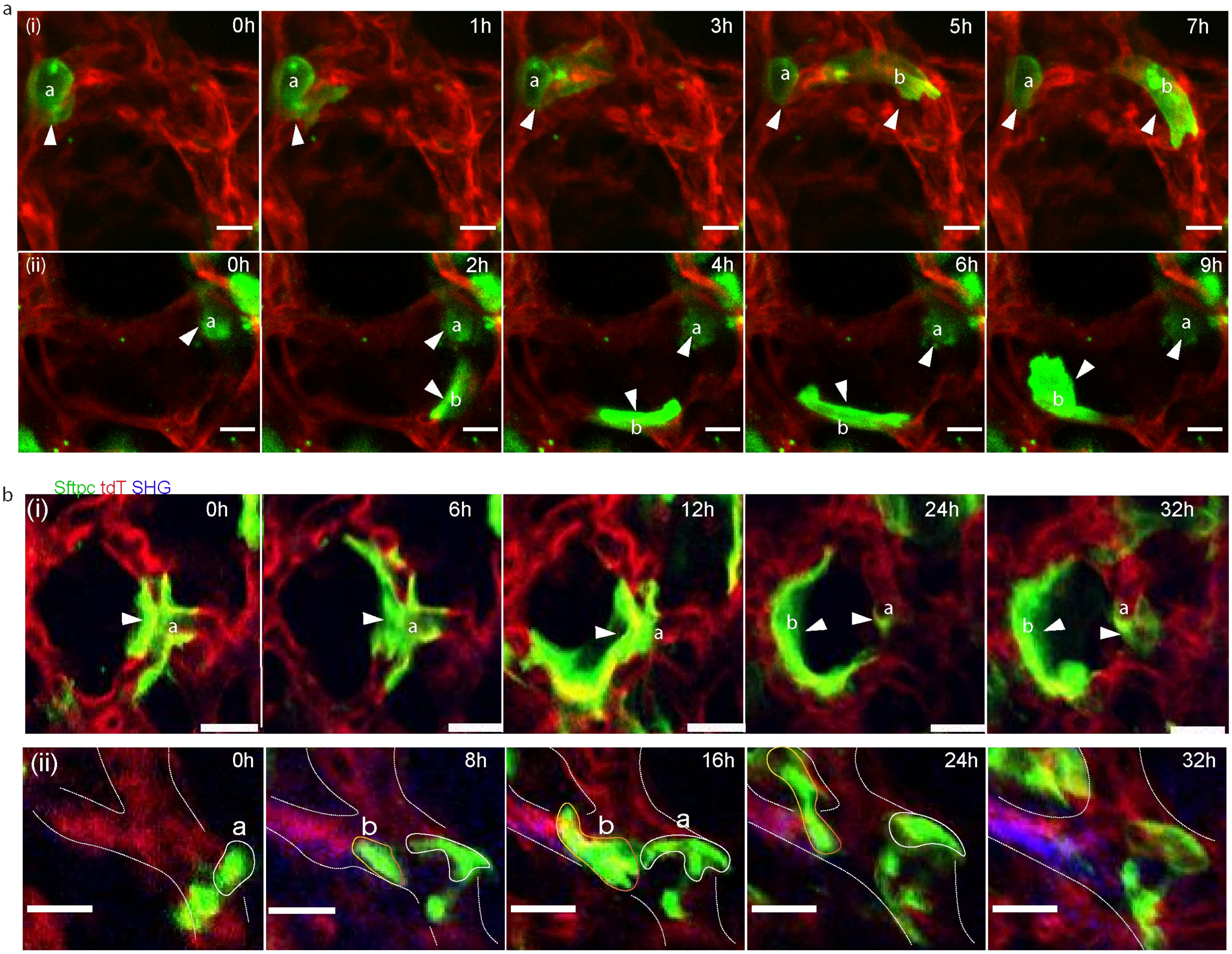

### Supp Figure 3

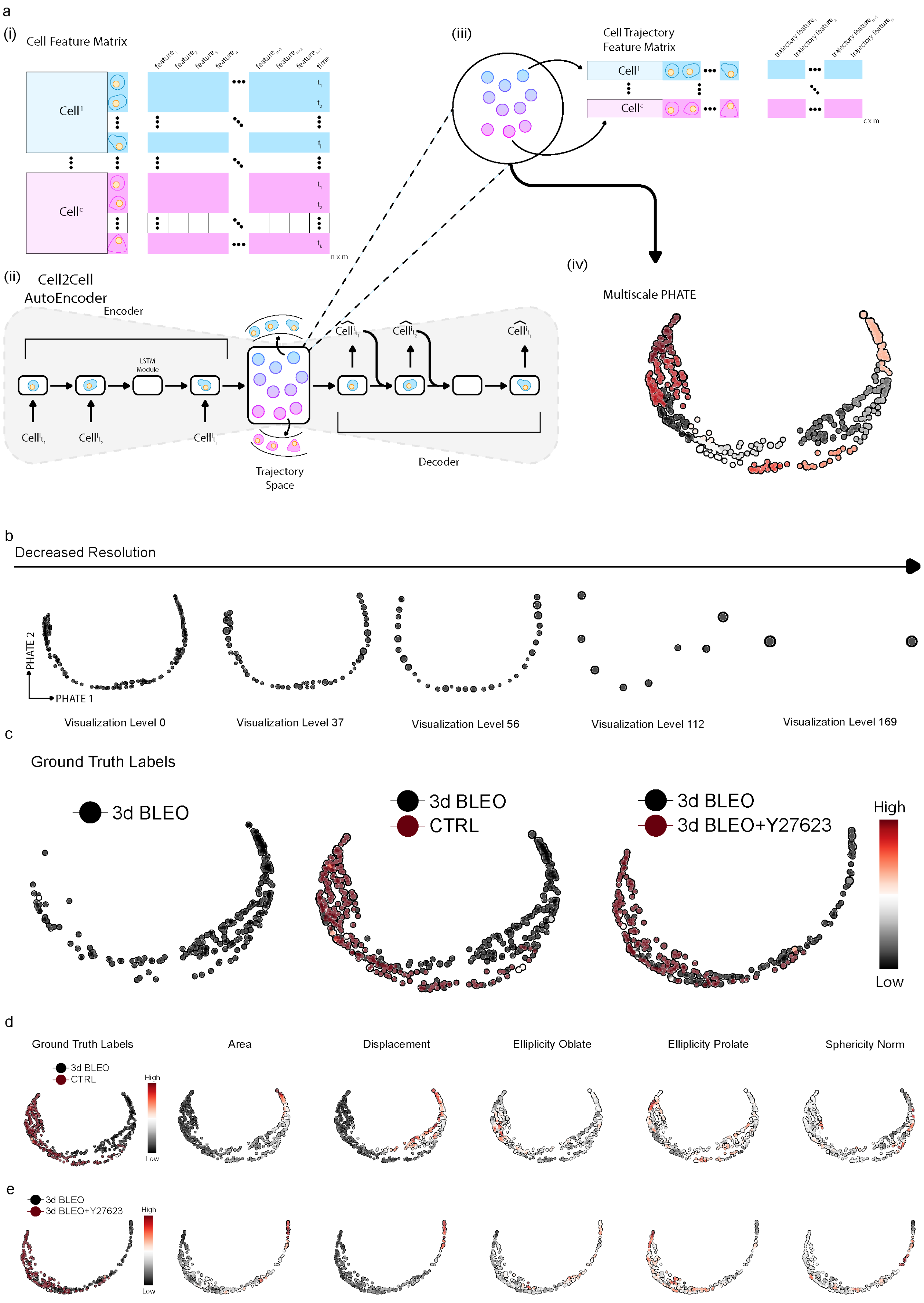

**SUPPLEMENTARY VIDEOS**

Supp Videos can be accessed at:
https://www.dropbox.com/scl/fo/1zrq3d1409yft2ic3yz81/h?dl=0&rlkey=2oh2qm09jmlaxz1zlnxano3rr

**Supp Video 1.avi** Ex-vivo time-lapse videos of fluorescent labelled AT2 cell (GFP+) 3 days post-bleomycin injury
**Supp Video 2.avi** Ex-vivo time-lapse videos of fluorescent labelled AT2 cell (GFP+) moving to the adjacent alveolus 3 days post-bleomycin injury
**Supp Video 3.avi** Ex-vivo time-lapse videos of fluorescent labelled AT2 cell (GFP+) 10 days post-bleomycin injury
**Supp Video 4.avi** In-vivo intravital timelapse imaging through permanent lung window of fluorescent labelled AT2 cell (GFP+) 3 days post-bleomycin injury
**Supp Video 5.avi** In-vivo intravital timelapse imaging through permanent lung window of fluorescent labelled AT2 cell (GFP+) 3 days post-bleomycin injury
**Supp Video 6.avi** Ex-vivo time-lapse videos of fluorescent labelled AT2 cell (GFP+) undergoing cell division 3 days post bleomycin injury.
**Supp Video 7.avi** Ex-vivo time-lapse videos of fluorescent labelled AT2 cell (GFP+) undergoing cell division 3 days post bleomycin injury.
**Supp Video 8.avi** In-vivo intravital timelapse imaging through permanent lung window of fluorescent labelled AT2 cell (GFP+) undergoing cell division 3 days post bleomycin injury.
**Supp Video 9a.avi** Ex-vivo time-lapse videos of rendered fluorescent labelled AT2 cell (GFP+) moving along alveolar septum 3 days post bleomycin injury
**Supp Video 9b.avi** Ex-vivo time-lapse videos of rendered fluorescent labelled AT2 cell (GFP+) moving along alveolar septum 3 days post bleomycin injury (from a different perspective).
**Supp Video 10.avi** Ex-vivo time-lapse videos of fluorescent labelled AT2 cell (GFP+) moving to the adjacent alveolus 3 days post bleomycin injury
**Supp Video 11.avi** In-vivo intravital timelapse imaging through permanent lung window of fluorescent labelled AT2 cell (GFP+) moving to the adjacent alveolus 3 days post bleomycin injury.
**Supp Video 12.avi** Ex-vivo time-lapse videos of fluorescent labelled AT2 cell (GFP+) elongating 3 days post bleomycin injury
**Supp Video 13.avi** Ex-vivo time-lapse videos of fluorescent labelled AT2 cell (GFP+) moving and enlarging 3 days post bleomycin injury
**Supp Video 14.avi** Ex-vivo time-lapse videos of 2 fluorescent labelled AT2 cells (GFP+) moving and enlarging 3 days post bleomycin injury
**Supp Video 15.avi** In-vivo intravital timelapse imaging through permanent lung window of fluorescent labelled AT2 cell (GFP+) moving and enlarging 3 days post bleomycin injury
**Supp Video 16.avi** In-vivo intravital timelapse imaging through permanent lung window of fluorescent labelled AT2 cell (GFP+) moving and elongating 3 days post bleomycin injury
